# An autophagy adaptor TRIAD3A promotes tau fibrillation by phase separation

**DOI:** 10.1101/2023.09.28.559893

**Authors:** Jiechao Zhou, Yang ‘an Chuang, Javier Redding-Ochoa, Alexander J. Platero, Alexander H. Barrett, Juan C. Troncoso, Paul F. Worley, Wenchi Zhang

## Abstract

Multiple neurodegenerative diseases are characterized by aberrant proteinaceous accumulations of tau. Here, we report an RBR-type E3 ligase TRIAD3A functions as a novel autophagy adaptor for tau. *TRIAD3A*(RNF216) is an essential gene with mutations causing age-progressive neurodegeneration. Our studies reveal that TRIAD3A E3 ligase catalyzes a novel mixed K11/K63 polyubiquitin chain and self assembles into liquid-liquid phase separated (LLPS) droplets. Tau is ubiquitinated and accumulates within TRIAD3A LLPS droplets and via LC3 interacting regions targets tau for autophagic degradation. Unexpectedly, tau sequestered within TRIAD3A droplets rapidly converts to amyloid aggregates without the transitional liquid phase of tau. In vivo studies reveal TRIAD3A decreases the accumulation of phosphorylated tau in a tauopathy mouse model, and disease-associated mutation of TRIAD3A increases accumulation of phosphorylated tau, exacerbates gliosis, and increases pathological tau spreading. In human Alzheimer’s disease brain, TRIAD3A colocalizes with tau amyloid in multiple histological forms suggesting a role in tau homeostasis. TRIAD3A is the first autophagic adaptor that utilizes E3-ligase and LLPS as a mechanism to capture cargo and appears especially relevant to neurodegenerative diseases.

## Introduction

Tauopathies include multiple forms of progressive neurodegenerative diseases and represent the leading cause of dementia. They are characterized by deposits of tau amyloid filaments ^1, 2^, which are usually accompanied by abnormal accumulation of other protein aggregates^3, 4^, such as amyloid-β, α-synuclein, TDP-43, and recently recognized TMEM106B ^5-7^. Sufficient amounts of evidence demonstrate tau proteostasis can be regulated by clearance of tau insoluble aggregates via aggrephagy receptors P62^8, 9^ and CCT2^10^, or by dissolving tau aggregates via VCP^11^ and TRIM11^12^. While emerging evidence indicates assembly of tau into fibrillar aggregates typically requires liquid-to-solid phase transition of tau liquid condensates induced under various supraphysiological experimental conditions like the addition of anionic co-factors or using truncated proteins ^13-17^, comparatively little is known about what drives the initiation of tau aggregation *in vivo*.

TRIAD3A(RNF216) is an E3 ubiquitin ligase of the ‘RING-in-between-RING’ (RBR) class, which harbors three tandem zinc-binding domains termed RING1, in-between RING (IBR) and RING2 (collectively called an RBR domain) (Figure 1a)^18^. Missense mutations in human gene TRIAD3A cause adult-onset neurodegenerative disorders including Huntington-like diseases (HDL) and Gordon-Holmes syndrome (GHS), the latter characterized by neuronal degeneration with cortical and cerebellar atrophy and age-progressive dementia^19-23^. Recent studies provide conflicting evidence regarding the polyubiquitin chain specificity assembled by TRIAD3A^24-27^, Therefore it remains to be clarified how TRIAD3A determines the fate of its substrates.

**Figure 1.**
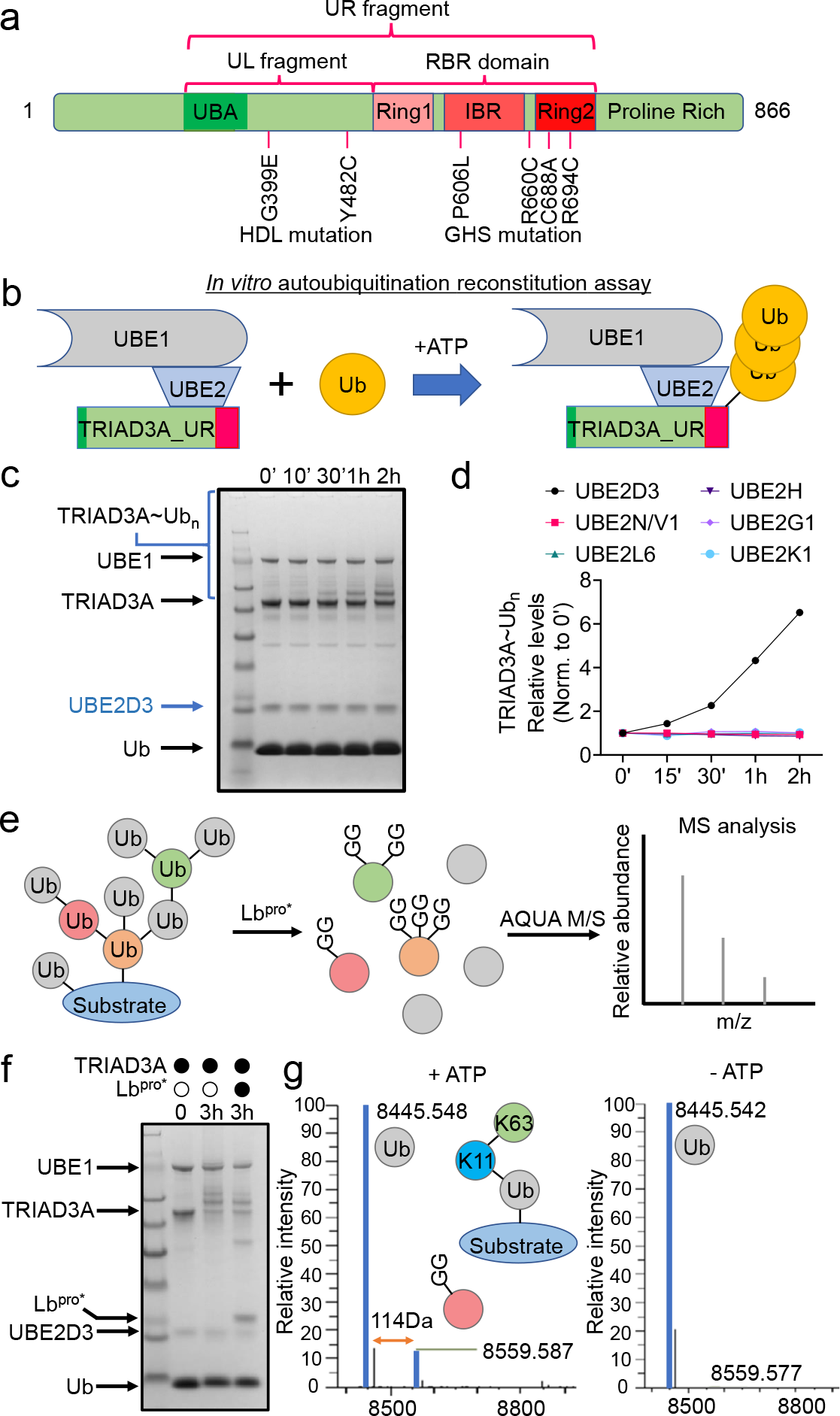
TRIAD3A assembles mixed K11/K63 linkage Ub chain. (a) Schematic representation of the *in silico*-prediction of domain organization of human TRIAD3A and selected point mutations identified in neurological patients. Ubiquitin-associated (UBA) domain and Ring1-IBR-Ring2 (RBR) domain are colored as indicated. HDL: Huntington-like diseases; GHS: Gordon-Holmes syndrome. (b) Cartoon depicting core components of TRIAD3A *in vitro* autoubiquitination reconstitution assay. (c) Identification of E2 enzymes that form ubiquitination modification on TRIAD3A. SDS-PAGE gels of time-course assay showing TRIAD3A autoubiquitination with UBE2D3. (d) Quantification results of time course measuring ubiquitination modified TRIAD3A for several selected UBE2s (Extended data Figure 1a, b and c). (e) Schematic of the Ub-clipping intact MS protocol that Lb^pro^* generates GlyGly-modified ubiquitin species. (f) SDS-PAGE gel image representing TRIAD3A autoubiquitination assay before and after Lb^pro^* treatment for 3 hours. (g) TRIAD3A assembles mixed K11/K63 polyUb. Quantification by spectra deconvolution of individual ubiquitin species from (f) with or without addition of ATP in autoubiquitination assay.

In the current work, we examine the polyubiquitin chain topology formed by TRIAD3A and identify TRIAD3A as a novel adaptor for autophagy. TRIAD3A undergoes E3 ligase-dependent phase separation to form liquid-like droplets regulated by specific polyubiquitin chains, and encodes LC3-interacting motifs that mediate its association with autophagosomes. A nonbiased screen for substrates using *in vivo* expressed TRIAD3A linked to Turbo peroxidase identifies multiple candidates linked to neurodegenerative disease. We confirm that TRIAD3A ubiquitinates tau and converts tau to amyloid solid aggregates in TRIAD3A mediated phase separation. Consistent with this TRIAD3A-tau-autophagy pathway, inhibition of lysosomes increases accumulation of tau amyloid. Furthermore, *in vivo* evidence presented here, including in human AD brain tissue, indicate that TRIAD3A initiated phase separation which underlies a novel tau aggregation pathway, confines tau and facilitates its autophagic turnover, which is disrupted in AD. Findings present the first example of physiological tau fibrillation and suggest a role for selective TRIAD3A mediated autophagy in the pathogenesis of tau fibrillation and neurodegenerative disease.

## Result

### TRIAD3A assembles mixed K11/K63 linkage Ub chains

RBRs bind E2 enzymes and serve as the ubiquitin-carrying domain. Ubiquitin (Ub) is transferred from E2 to a catalytic cysteine of RING2, and subsequently to the substrate or an acceptor Ub to form polyubiquitin (polyUb) chains^18, 28^. To investigate polyUb chain linkage specificity of E3 ligase TRIAD3A, we developed an *in vitro* assay and monitored autoubiquitination (Figure 1b). TRIAD3A_UR (UR fragment: amino acid 260-750; Figure 1a) was used to screen a panel of different E2 enzymes^29^. We found UBE2D3 exhibits significant *in vitro* ligase activity, compared with other E2s (Figure 1c and Extended data Figure 1a, b and c). Mass spectrometry revealed K11 and K63 linkages to Ub (Extended data Figure 1d, e). To further assess polyUb chain architecture, we used Ub-clipping and intact ubiquitin mass analysis^30^ (Figure 1e) and observed exclusively mono-GG modified ubiquitin species indicating an absence of branched polyubiquitin (Figure 1f and g). These data establish an *in vitro* TRIAD3A autoubiquitination assay and demonstrate TRIAD3A assembles novel mixed K11/K63 linkage Ub chains.

### PolyUb facilitates TRIAD3A ligase activity

In silico analysis of a conserved region at the N-terminus of TRIAD3A RBR domain identified a potential ubiquitin associated domain (UBA)^31, 32^ (Figure 2a). To evaluate its ubiquitin-binding properties, GST pull-down experiments were performed with TRIAD3A_UL fragment construct (UL fragment: amino acid: 260-509; Figure 1a). As shown in Figure 2b, TRIAD3A_UL preferentially bound diUb over monoUb, and displayed no selectivity for different ubiquitin chain linkages. We compared the predicted 3D structure of TRIAD3A UBA domain with UBA domains complexed with diUb or monoUb in the PDB library (Extended data Figure 2a and b) and identified a conservative Ub interaction surface (Figure 2c). Point mutations R294A and L323A of the putative UBA domain disrupted binding to diUb (Figure 1d). To further test the potential regulatory role of UBA domain in TRIAD3A E3 ligase activity, we added various polyUbs of different chain lengths or chain specificities to TRIAD3A *in vitro* autoubiquitination reconstitution reactions. TriUb and tetraUb enhanced TRIAD3A ligase activity but not the diUb (Figure 1e and f; Extended data Figure 2c). This action was not specific for polyUb chain linkage-type. We further used our assay with addition of polyUb to determine that mutations in RBR domain identified in GHS (Extended data Figure 2d) abolish chain formation reaction (Figure 1g and h). Importantly, L323A mutation prevented polyUb-mediated increase of TRIAD3A ligase activity (Figure 1g and h). These data indicate that TRIAD3A E3 activity can be regulated by local polyubiquitin via its UBA domain.

**Figure 2.**
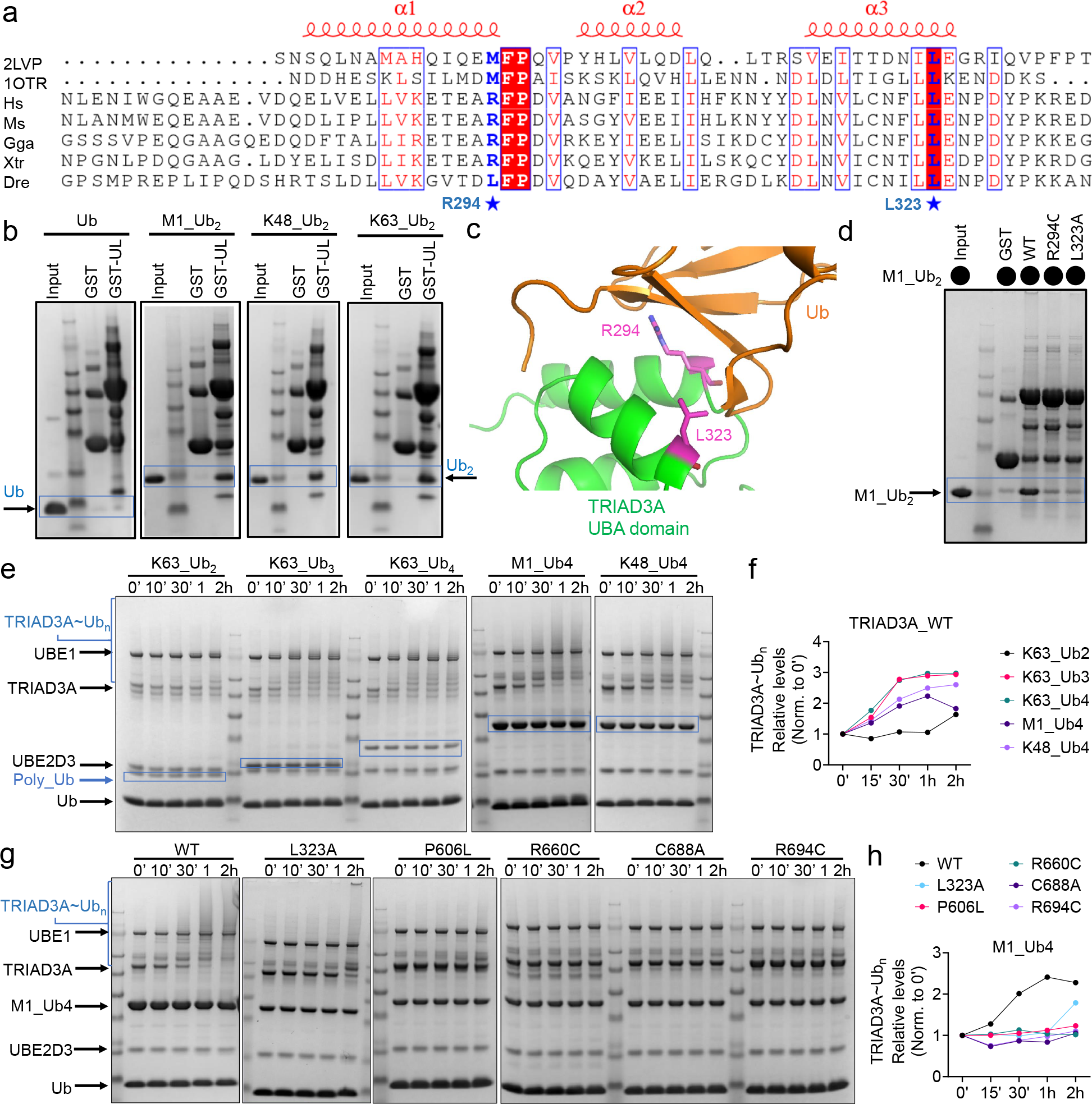
PolyUb facilitates TRIAD3A E3 ligase activity. (a) The sequence alignment of TRIAD3A UBA domain for typical species based on its 3D structural superposition with UBA domains complexed with Ub (PDB ID: 1OTR) and diUb (PDB ID: 2LVP). The structures of human TRIAD3A UBA domain was predicted by DeepMind with AlphaFold. Secondary structures of TRIAD3A UBA domain are shown on the top in red. Conservative residues of the UBA domain making direct contact with ubiquitin are marked with a blue star on the bottom. (b) SDS-PAGE gels of *in vitro* GST pull-down experiments testing the binding of WT TRIAD3A UL region (aa 260-509) to different types of diUb. (c) Magnified view of the conservative interaction interface of predicted TRIAD3A UBA domain in green with ubiquitin in brown. The residues at the interaction interface are shown as a stick representation in purple. (d) Mutations in the UBA domain (R294A and L323A) at the binding interface disrupted the interaction with diUb determined by *in vitro* GST pull-down experiment. (e) SDS-PAGE gels of *in vitro* TRIAD3A autoubiquitination assays showing different types of polyUb with longer chain than diUb enhance TRIAD3A *in vitro* ligase activity. (f) The graph shows quantification of time course in (e) measuring ubiquitination modified TRIAD3A with addition of polyUb via densitometry. (g) SDS-PAGE gel images of TRIAD3A ubiquitination assay showing that mutation of Ub interacting residue in the UBA domain prevents TRIAD3A activity increase in presence of M1_Ub_4_, and missense mutations in the RBR domain identified in neurological disorders disrupt TRIAD3A ligase activity with addition of M1_Ub_4_. (h) The graph shows quantification of time course in (g) measuring ubiquitination modified TRIAD3A for mutations in UBA domain or RBR domain with addition of M1_Ub_4_ via densitometry.

### TRIAD3A forms E3 ligase activity-dependent droplets

We noted that although reconstitution reactions included excess ubiquitin and TRIAD3A, the rate of TRIAD3A autoubiquitination decreased over time (Figure 1e), suggesting TRIAD3A shifted to a slow dynamic state. We considered that TRIAD3A can be modified by polyUb that may interact with its UBA domain. Since network interactions among macromolecules can drive phase separation^33, 34^, we further hypothesized that interaction between polyubiquitin chains on TRIAD3A and the UBA domain may promote phase separation that slows TRIAD3A autoubiquitination (Figure 3a). To test this hypothesis, we used recombinant mGFP-TRIAD3A_UR fragment (260-750), which includes the UBA and RBR domains, to reconstitute autoubiquitination reaction. We observed phase separation without artificial crowding agents (Figure 3a to e; Extended data Figure 3a and b). In the control reaction without adenosine triphosphate (ATP), TRIAD3A phase separation did not occur (Extended data Figure 3a and b). ATP together with M1_Ub4 promoted TRIAD3A condensate formation (Extended data Figure 3a and b). Measures of sphericity, recovery of fluorescence after photobleaching (Figure 3b), and observation of droplets formation (supplemental video 1 and 2) indicated that condensates are liquid-like droplets. To determine the effect of polyUb chain specificity, we added various polyUbs of different chain lengths or chain specificities to mGFP-TRIAD3A_UR *in vitro* autoubiquitination reconstitution reaction. Multiple forms of polyUbs promote TRIAD3A phase separation including K63_Ub2, linear ubiquitin tetramer (M1_Ub_4_), K48_Ub_4_ and mixed K48/K63_Ub_4_ (Figure 3c and d). Point mutations of the UBA (L323A) and RBR (R694C) domains confirmed that TRIAD3A driven phase separation requires UBA binding to Ub and E3 ligase activity (Figure 3e). Interestingly, K63_Ub_3_ and K63_Ub_4_, which do bind the UBA and induce E3 ligase do not induce TRIAD3A phase separation (Figure 2e and f, Figure 3c and d). These findings indicate that the UBA binding and E3 ligase activity alone are not sufficient to determine phase separation and suggest regulatory specificity of polyubiquitin within the liquid-like droplets of TRIAD3A. In line with *in vitro* assays, full-length mCherry-TRIAD3A expressed in N2A cells exhibited condensates with sphericity, recovery of fluorescence after photobleaching (Extended data Figure 3c, d and f), and fusion of TRIAD3A condensates (Extended data Figure 3e). In contrast to *in vitro* assays, a predominant fraction (32/40) of TRIAD3A condensates in N2A cells appeared to be associated with organelles (ex. autophagosome, as below) and did not display liquid-like properties (Extended data Figure 3c, bottom). Collectively, these data demonstrate that TRIAD3A undergoes phase separation to form liquid droplets and that E3 ligase activity and liquid phase separation can be independently regulated by polyubiquitin chain linkage.

**Figure 3.**
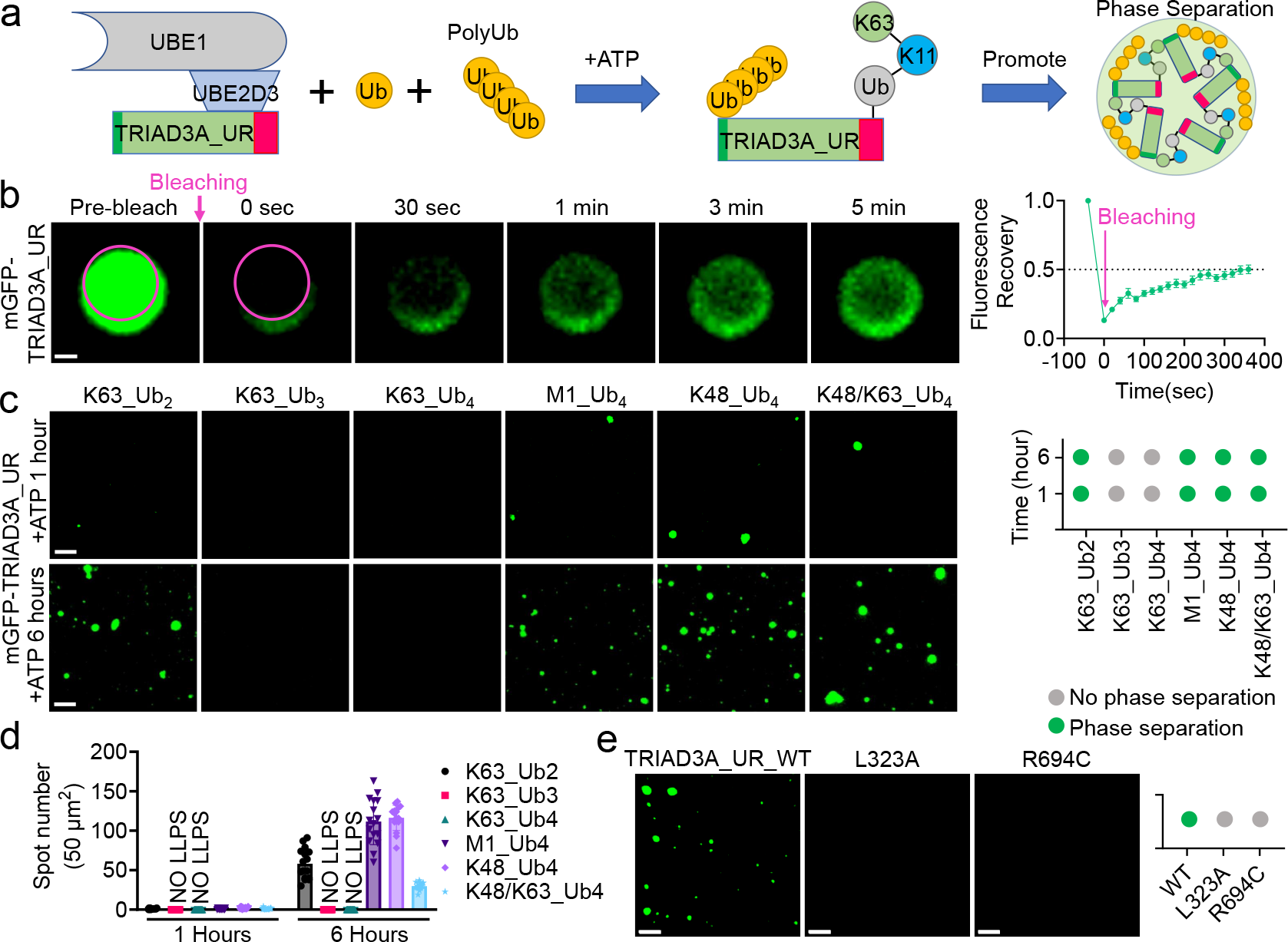
PolyUb induces TRIAD3A phase separation. (a) Schematic illustration of TRIAD3A activity induced phase separation mediated by network interaction of UBA domain with polyUb on TRIAD3A. (b) Left: fluorescence intensity recovery of a TRIAD3A droplet after photobleaching. Scale bar, 0.5 μm. Right: Quantification of fluorescence intensity recovery of photobleached TRIAD3A droplets. (c) Left: TRIAD3A droplets formation in presence of different type of polyUb at two time points: 1 hour and 6 hours. Scale bar, 5 μm. Right: Phase diagram of the formation of TRIAD3A droplets at indicated time points with different polyUb types. (d) The spot number of TRIAD3A droplets were quantified in the presence of different type of polyUb. Each spot represent one imaged area. (e) Formation of TRIAD3A droplets and their inhibition by mutations in the UBA domain (L323A) and RBR domain (R694C) in presence of M1_Ub_4_. Scale bar, 5 μm.

### TRIAD3A is an autophagy adaptor

Liquid-like condensates play important roles in triaging protein cargos for selective autophagic degradation^35, 36^ via autophagy adaptors that simultaneously contain ubiquitin-binding domain and LC3 interaction regions (LIR) ^37, 38^. In the disordered N-terminal region of TRIAD3A we identified two consensus LIR motifs that are conserved in mammals (Figure 4a and b). We applied AlphaFold2 ^39^ to generate complex structures of LC3B with TRIAD3A LIRs (Figure 4c) revealing two hydrophobic residues that contact two hydrophobic pockets of LC3B^40^. In vitro GST pulldown experiments confirmed the physical interaction of LC3B and TRIAD3A, while point mutations in TRIAD3A LIR motifs reduced LC3B binding (Figure 4d). TRIAD3A transgene in N2A cells interacted with GFP-LC3B, whereas the double-point mutation W24A/F77A in LIR region, R694C in RBR domain, or deletion of N-terminus that contains the LIR region (delete amino acid 1-183), attenuated their interaction (Extended data Figure 4a). In N2A cells TRIAD3A transgene co-localized with GFP-LC3B and endogenous autophagosome membrane marker WIPI2^41,42^(Extended data Figure 4b and c). Their co-localized puncta became larger and more numerous following inhibition of lysosome function by treatment with bafilomycin A1 (Extended data Figure 4b and c). The ability of TRIAD3A to sequester to LC3B-positive autophagosomes was abolished by TRIAD3A mutations of LIR(W24A_F77A) or RBR(R694C) domains (Extended data Figure 4b and c). Examining native proteins in primary mouse astrocytes treatment with Bafilomycin A1 or Hydroxychloroquine induced TRIAD3A to form puncta with LC3B (Figure 4e and f). Together, these data indicate that TRIAD3A associates with the autophagy machinery and functions as a novel autophagy adaptor (Figure 4g).

**Figure 4.**
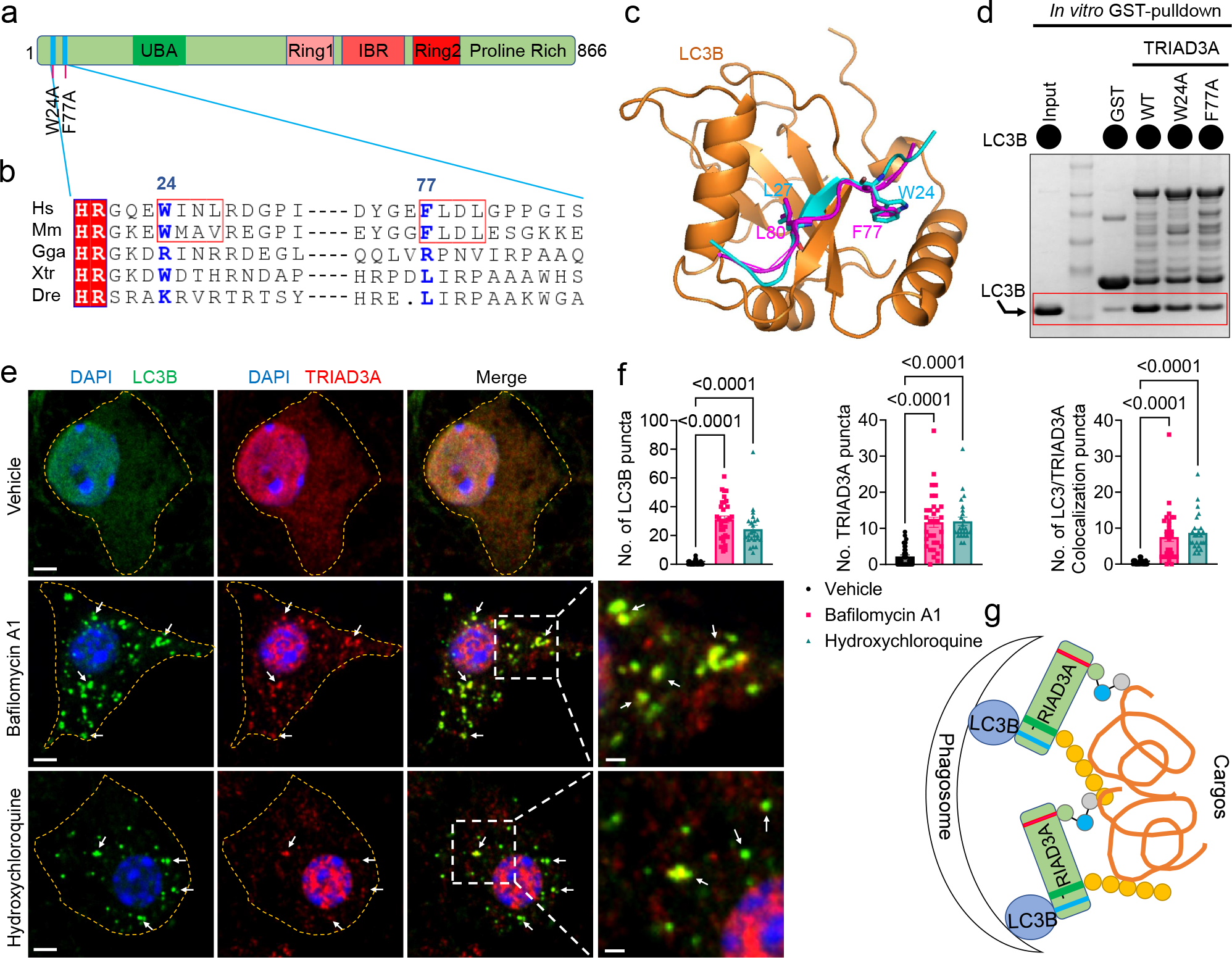
TRIAD3A is an autophagy adaptor. (a) Position of LC3 interaction region (LIR) in the full-length of human TRIAD3A. (b) The sequence alignment of TRIAD3A N-terminal disordered region for typical species. Two LIRs are marked with a red box and conservative in mammals. (c) Superposition of complex structures of two TRIAD3A LIR fragments (aa 21-31 in cyan or aa 75-84 in purple) with LC3B (brown). This complex structure is generated by AlphaFold2, respectively. (d) SDS-PAGE gel of *in vitro* GST pull-down experiment showing mutations in the TRIAD3A LIRs reduce binding to LC3B. (e) Representative immunofluorescence images of primary cultured astrocytes with TRIAD3A and LC3B antibodies. The cells were treated with Bafilomycin A1 (10nM, 24 hours) and hydroxychloroquine (10μM, 24 hours). White arrows indicated the TRIAD3A colocalized with LC3B. Scale bar, 3 μm. Scale bar of magnification images, 1 μm. (f) Quantification data of LC3B puncta, TRIAD3A puncta and LC3B colocalized with TRIAD3A puncta. Each spot represents per cell, vehicle, N = 38; Bafilomycin A1, N = 32; hydroxychloroquine, N = 25. p value was determined by one-way ANOVA. (g) Schematic illustration of TRIAD3A as an autophagy adaptor. Data are presented as mean ± SEM.

### TRIAD3A induces fibrillar tau aggregates in droplets

To identify cargo content involved in TRIAD3A-mediated selective autophagy, we expressed TRIAD3A fused to an engineered biotin ligase (TurboID) by viral transduction in mouse brain neurons (AAV-CAMKII-TRIAD3A-V5-TurboID). After treatment with biotin substrate, brain was harvested and biotinylated proteins were identified by liquid chromatography-tandem mass spectrometry^43^ (Extended data Figure 5a to f). Top candidates included proteins linked to neurodegenerative diseases: tau, hnRNPs, TARDBP/TDP-43, Fus^44^ (Extended data Figure 5d and f). We selected tau for further analysis. Immunoprecipitation (IP) assays from TauP301S mice brains confirmed that endogenous TRIAD3A interacts with human tau (Figure 5a). In vitro GST-pulldown experiments demonstrated TRIAD3A_UR binds recombinant human tau 0N4R. TRIAD3A_UR and UL fragments preferentially bind tau repeat region 3 (Tau_R3) compared to tau repeat region 4 (Tau_R4) (Extended data Figure 5f and h). To assess if TRIAD3A can ubiquitinate tau, we monitored recombinant human tau 0N4R conjugated with Alexa Fluor 488 maleimide to distinguish it from similar sized TRIAD3A_UR and confirmed tau is ubiquitinated by TRIAD3A *in vitro* (Figure 5b). TRIAD3A also promotes tau ubiquitination in HEK293 cells (Figure 5c) while point mutations that impair TRIAD3A liquid-like condensate formation or association with LC3B attenuate tau ubiquitination *in vivo* (Figure 5c). Importantly, tau 0N4R coacervates (co-partitions) into TRIAD3A condensates in the *in vitro* autoubiquitination reconstitution assay. Unlike TRIAD3A, tau clusters in TRIAD3A droplets were not uniformly distributed (Figure 5d, e and f) and did not recover after photobleaching (Figure 5h). We performed a droplet sedimentation assay with unlabeled tau to confirm TRIAD3A droplets enrich ubiquitinated tau (Figure 5g). Moreover, tau within TRIAD3A droplets stained with thioflavin S (ThioS) indicating fibrillization (Figure 5i and j). These data indicate TRIAD3A induces tau fibrillization in association with phase separation and ubiquitination in TRIAD3A droplets. We refer to the conversion process of tau as “nested phase separation” (Figure 5k) to highlight the role of TRIAD3A in leading to tau amyloid solid aggregates.

**Figure 5.**
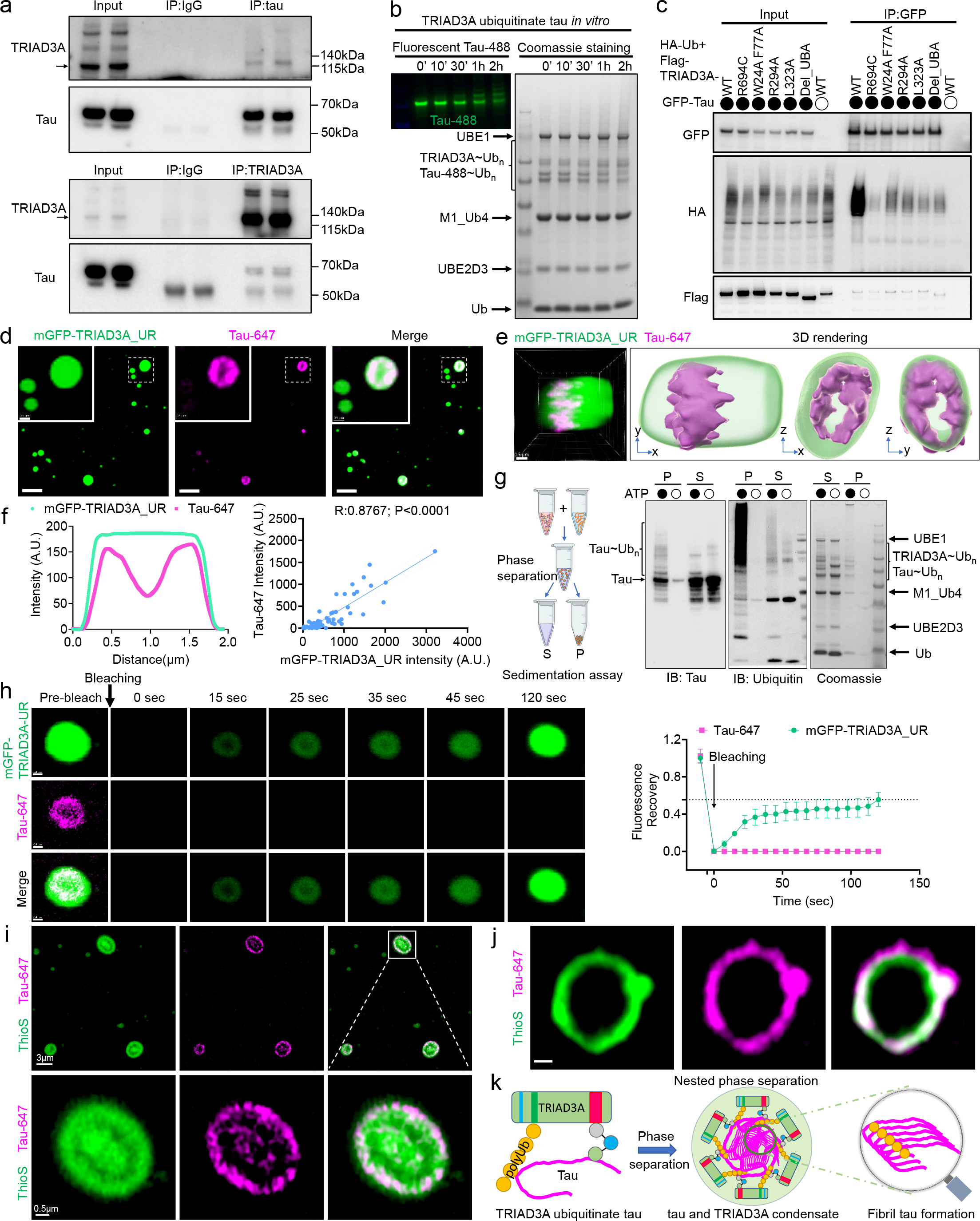
TRIAD3A promotes tau amyloid aggregation in phase separation droplets (“nested phase separation”). (a) Representative western blot data of TRIAD3A and tau CoIP. CoIP of TRIAD3A and tau from 9-month-old TauP301S mice brains using TRIAD3A antibody and tau(A10) antibody. Immunoblot with TRIAD3A and tau antibodies showed their endogenous interaction. (b) Representative images of TRIAD3A ubiquitination reaction with addition of tau. Left: Tau ubiquitination was monitored by fluorescently labeled full-length recombinant 0N4R tau with Alexa Fluor 488 (Tau-488). Right: SDS-PAGE gel of time-course assay showing TRIAD3A autoubiquitination and tau ubiquitination. (c) GFP-Tau and HA-Ub were transfected into HEK293 cells together with wildtype TRIAD3A or indicated mutations in various domains. Immunoprecipitated of GFP antibody from the cell lysates. Representative western blot images shown of ubiquitination of GFP-Tau by various TRIAD3A. GFP, HA and Flag were analyzed by western blotting. (d) Co-occurrence of TRIAD3A and tau in the droplets during TRIAD3A autoubiquitination reaction with addition of Alexa Fluor 647 labeled full-length recombinant 0N4R tau (Tau-647). Scale bar, 3 μm. Inset: 0.5 μm. (e) Three-dimensional (3D) rendering of Tau-647 in TRIAD3A droplet shows ununiform distribution of tau. Scale bar, 0.5 μm . (f) left panel: Radial distribution of the fluorescence intensities of TRIAD3A and Tau-647 in the droplet indicated that the distribution of tau was ununiform. Right panel: correlation analysis of the fluorescence intensities of TRIAD3A and Tau-647. p value was determined by two-tailed t test. (e) left panel: Schematic diagram of the droplets sedimentation assay to separate the condensed liquid droplets and the supernatant by centrifugation. Right panel: The pellet and supernatant of TRIAD3A droplets formed autoubiquitination reaction in presence of full-length recombinant 0N4R tau were analyzed by Coomassie staining and western blot using anti-tau and anti-ubiquitin antibody. S, supernatant; P, pellet. (h) Left: Representative image of fluorescence intensity recovery of a TRIAD3A and Tau-647 droplet after photobleaching, but no recovery of Tau-647. Scale bar, 0.4 μm. Right: quantification of fluorescence intensity recovery of photobleached TRIAD3A and Tau-647 droplets. (i and j) Representative images of Thioflavin S (ThioS, a amyloid cross-β-sheet binding dye that labels tau filaments) staining showed strong co-localization and co-distribution with Tau-647. Panel I scale bar 3 μm; inset 0.5 μm. Panel J Scale bar, 0.4 μm. (k) Schematic illustration of ‘nested’ tau phase separation in TRIAD3A droplet.

### TRIAD3A regulates tau spreading and turnover

We asked whether TRIAD3A plays a role in turnover of tau. We co-expressed aggregation-prone tau mutant P301L with TRIAD3A in N2A cells and analyzed co-localization with LC3B. Upon digitonin treatment to remove soluble tau, we observed TRIAD3A, Tau_P301L, LC3B in puncta colocalizing with Amylo-Glo, a fluorescent dye for amyloid aggregates^45^ (Extended data Figure 6a). In addition, upon digitonin pretreatment to extract cytosolic LC3B^46^, mature autophagosome membrane-bound lipidated LC3B-II co-localized with TRIAD3A (Extended data Figure 6a). Consistent with the role of lysosomes in degradation of autophagosome content^47^, inhibition of lysosome acidification by Bafilomycin A1 lead to the accumulation of amyloid tau puncta visualized by Amylo-Glo (Extended data Figure 6a and b). By contrast, TRIAD3A_R694C mutation showed markedly decreased amyloid tau aggregates (Extended data Figure 6a and b).To determine if TRIAD3A acts *in vivo* to mitigate tau pathology, we monitored tau hyperphosphorylation in hippocampus and cortex of TauP301S transgenic mice following injections of AAV-CAMKII-V5-TRIAD3A (Extended data Figure 6c and d). TRIAD3A reduced phosphorylated tau detected by AT8 (p-S202/T205) antibody, a tau form linked to neurofibrillary tangle formation^1, 2^ (Figure 6a and b). Control experiments that expressed GFP did not reduce AT8 intensity (Figure 6a and b; Extended data Figure 6e to g). To strengthen our understanding of the role of TRIAD3A in tauopathy, we generated a mouse model carrying the disease-associated mutation R694C and examined brain of homozygous crosses with TauP301S mice. In 8-month-old mice, TRIAD3A(R694) mutation increased gliosis and tau aggregation (AT8 staining) (Figure 6c and d). We also examined an *in vivo* spreading model using AAV-GFP-P2A-hTauP301L^48^. This model distinguishes transduced neurons expressing both GFP and hTauP301L from neurons expressing only spread hTauP301L. Three months after injection of AAV-GFP-P2A-hTauP301L into hippocampus of WT and TRIAD3A(R694C) mutation mice (Figure 6e, f and g), we observed the anticipated spread of tau in the WT mice and prominently increased spread in the TRIAD3A(R694C) (Figure 6h and i). Increased spreading was not due to differences in the number of transduced cells, as a comparable intensity of GFP signal was observed in both groups (Figure 6h and i). These findings indicate that TRIAD3A loss of function can exacerbate pathological tau accumulation and propagation that contribute to abnormal tau homeostasis, which strongly implicate TRIAD3A in normal and pathological tau homeostasis.

**Figure 6.**
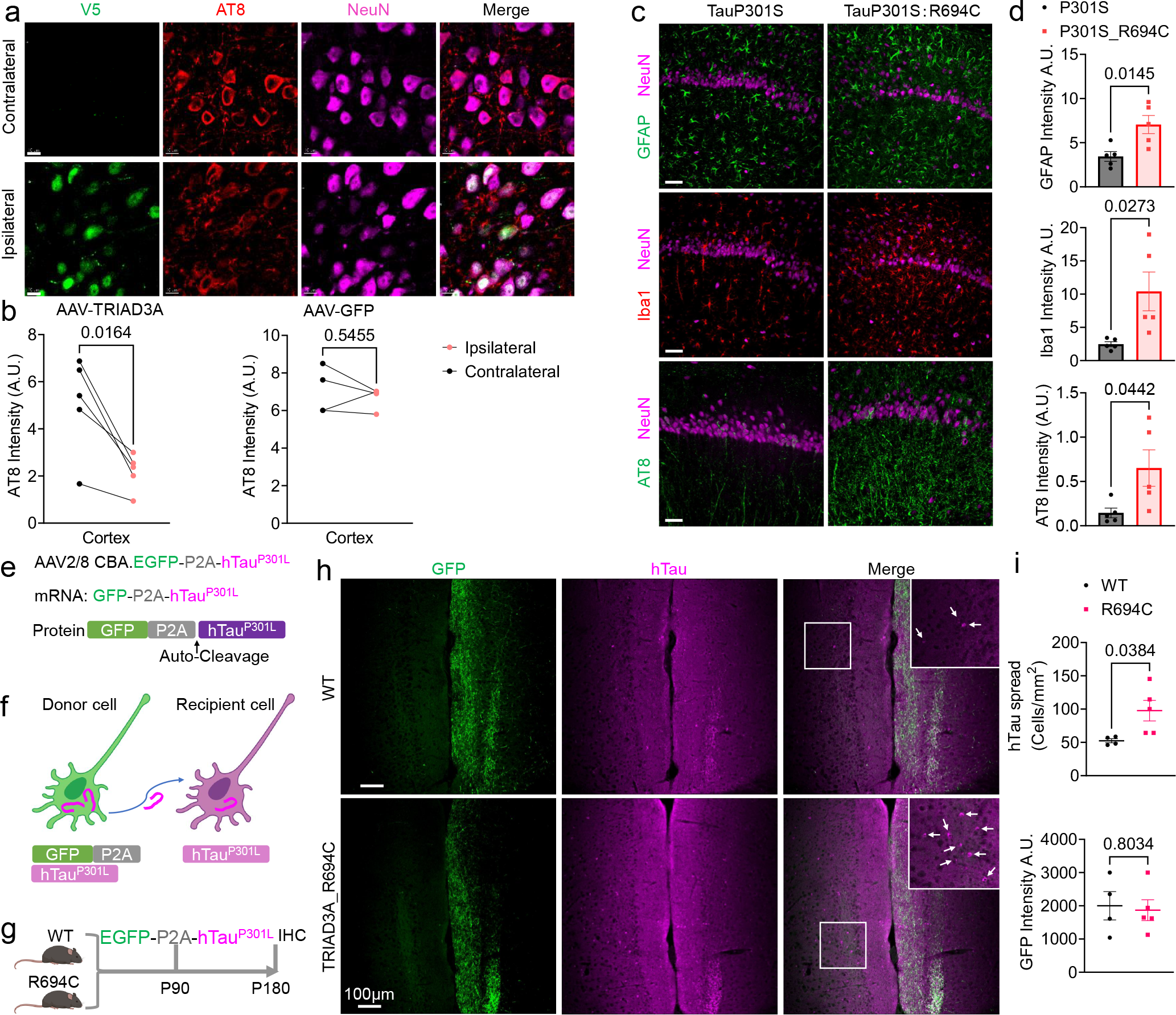
TRIAD3A modulates tau turnover. (a) Representative immunofluorescence images of NeuN (magenta), AT8 (red) and V5 (green) in cortex of AAV-TRIAD3A-V5 injected TauP301S mice brains. (b) Quantification of AT8 intensity in AAV-TRIAD3A (A) or AAV-GFP (Extended data Fig.6e) injected cortex of TauP301S mice brains. p value was determined by paired t test. Red: ipsilateral; black: contralateral. (c) Representative immunofluorescence images of Iba1/NeuN, GFPA/NeuN and AT8/NeuN in the hippocampus CA1 of 8-month-old TauP301S and TauP301S:TRIAD3A_R694C (TauP301S_R694C) mice. (d) Quantification of Iba1, GFAP and AT8 in the 8-months old mice. p value was determined by unpair t-test. TauP301S, n = 5; TauP301S_R694C, n = 5. Data are presented as mean ± SEM. (e) Schematic of the AAV-EGFP-P2A-hTauP301L construct. (f) Schematic of the AAV-EGFP-P2A-hTauP301L tau-spreading model in mouse brain. GFP positive neurons are hTau donors, spreading hTauP301L to recipient GFP negative neurons. (g) Timeline of injections in WT and R694C mice. Animals were injected with AAV-EGFP-P2A-hTauP301L at P90. Animals were euthanized at P180 to evaluate hTauP301L spreading in the mice brains. (h) Representative immunofluorescences images for hTau in WT and R694C mice overexpressing GFP-P2A-hTauP301L virus. Scale bar, 100 µm. White arrow indicated hTau positive cells in the cortex. (i) Quantification of cortical hTau positive/GFP negative cells (top) and GFP intensity (bottom) of injected mice. WT (n = 4) and R694C (n = 5), per mice have 3-5 sections for imaging. p value was determined by unpaired t-test. Data are presented as mean ± SEM.

### TRIAD3A in human brain and Alzheimer’s disease

To further investigate whether TRIAD3A is dysfunctional in human AD. We first examined TRIAD3A immunoreactivity (IR) in human brain, confirming localizations using two independent antibodies validated against TRIAD3A knockout mouse brain (Extended data Figure 7a and b). In normative aged control frontal cortex, TRIAD3A IR typically localizes to the soma and nucleus of neurons and astrocytes and stands out relative to prominent puncta in the surrounding neuropil (Figure 7a; Extended data Figure 7c). TRIAD3A IR is also present in occasional proximal dendrites. In AD brain, TRIAD3A IR appears generally reduced with fewer cells showing somatic staining and reduced density of puncta (Figure 7a; Extended data Figure 7c). TRIAD3A IR frequently co-localizes with structures staining for total tau (tau1 antibody) and Amylo-Glo suggesting association with tau amyloid (Figure 5a; Extended data Figure 7e to g). TRIAD3A IR also co-localized with p-S202/T205-Tau (AT8) at sites that co-localized with Amylo-Glo (Figure 7b and g; Extended data Figure 7c and d). Both double labeling strategies revealed associations with different forms of tau aggregates that represent stages of tau pathology including (from early to late) small puncta, fibrils, early tangles, and neurofibrillary tangles (Figure 7a). Consistent with a general reduction of TRIAD3A IR intensity TRIAD3A protein was reduced in western blot analysis of medial frontal and temporal cortex (MFC) of AD compared to age-matched controls (Figure 7c, d, e, f and g). These observations confirm the association of TRIAD3A with pathological forms of tau and suggest that in AD the TRIAD3A pathway maintains its partial function required to co-partition fibrillar tau but fails to effectively degrade tau.

**Figure 7.**
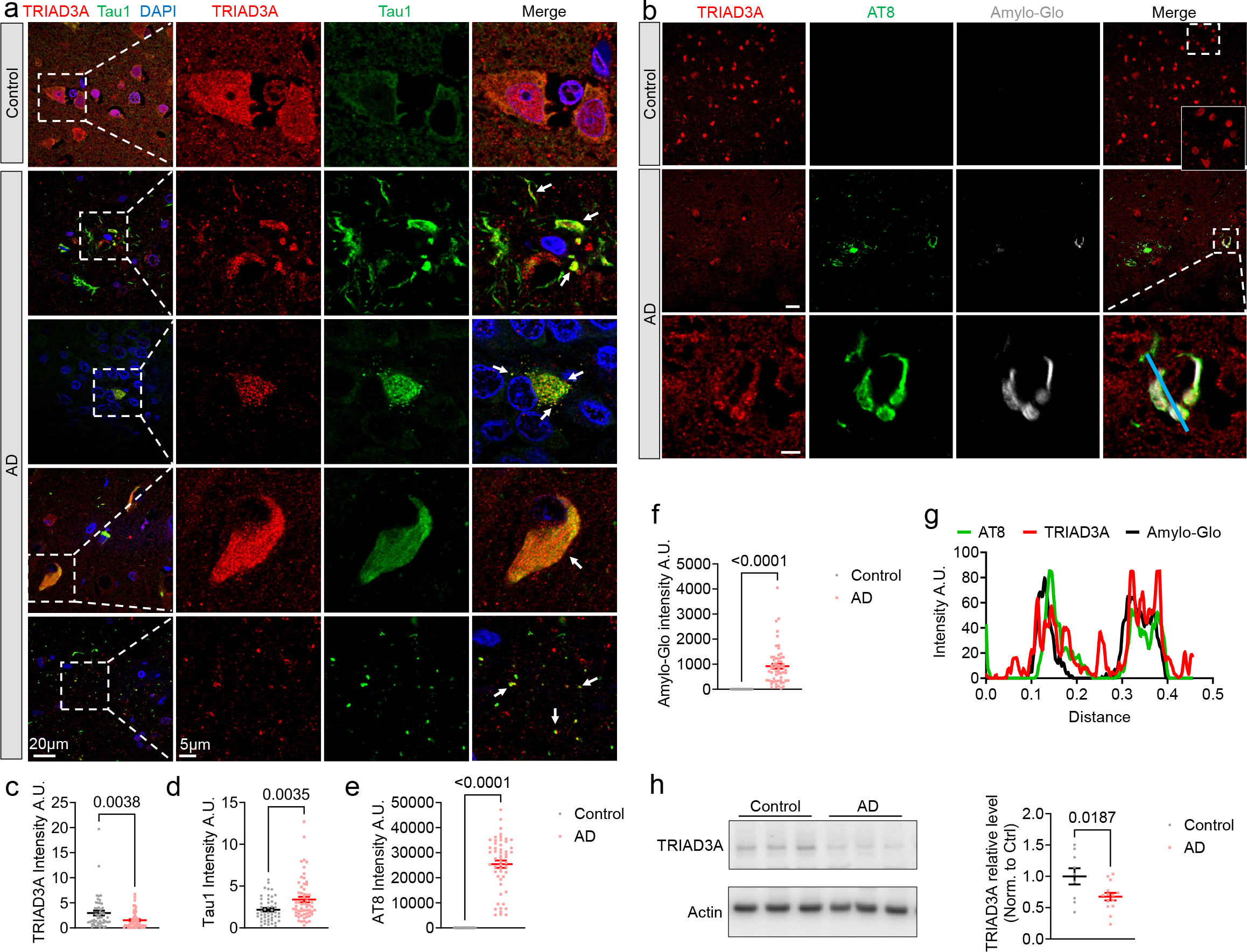
TRIAD3A dysregulated in Alzheimer’s disease. (a) Representative images of TRIAD3A and total-tau (tau1 antibody) in the human frontal cortex of nondementia (Control) and AD subjects. White boxes are magnified to the right. Scale bar, 20 μm. Magnified images scale bar, 5 μm. (b) Representative images of TRIAD3A and AT8 (anti-phospho-tau (Ser202, Thr205)) and Amylo-Glo in the frontal cortex of Control and AD subjects. White boxes are magnified to the bottom. Scale bar, 20 μm. Magnified images scale bar, 5 μm. (c, d, e, f) Quantification of TRIAD3A (b), tau1 (d), AT8 (e) and Amylo-Glo (f) intensity in Control and AD. Control, n = 50 images from 4-6 images/case, 2 sections/case and 5 control cases; AD, n = 60 images from 4-6 images/case, 2 sections/case and 5 AD cases. p value was determined by unpaired t test. Data are presented as mean ± SEM. (g) Radial distribution of the fluorescence intensities of AT8, TRIAD3A and Amylo-Glo indicated that TRIAD3A colocalization with AT8 and Amylo-Glo. (h) Representative western blot images shown TRIAD3A protein levels in the human Control and AD medium frontal cortex tissues. Quantification of TRIAD3A levels in Control and AD. Control, n = 9; AD, n = 14; p value was determined by unpaired t test. Data are presented as mean ± SEM.

## Discussion

The present study identifies TRIAD3A as a novel class of autophagy adaptor. TRIAD3A utilizes unique mixed K11/K63 polyUb assembly together with a UBA domain to generate liquid-like phase separated (LLPS) condensates, and LC3 interaction domains to target the complex to autophagosomes. TRIAD3A is distinct from other autophagic adaptors such as SQSTM1/p62 and NBR1, which lack E3 ligase activity and recruit protein aggregates linked to K63 polyUb^49^. Tau is a direct substrate of TRIAD3A and LLPS cargo exhibiting a striking conversion to a fibrillar state as its co-partitions or coacervates with TRIAD3A. This “nested phase separation” of tau occurs rapidly without artificial crowding agents, suggesting that autophagic processing of tau in normal physiological conditions utilizes a solid fibrillar intermediate in a manner mechanistically distinct from preferential autophagic clearance of the protein condensates with a certain amount of liquidity^50, 51^ and phase transition^16, 17^. Our observation unveils a novel aggregation pathway of tau from a soluble monomer into amyloid aggregates without the transitional liquid phase of tau. The aggregation-prone property of tau may be exploited to concentrate and confine the protein for autophagic processing.

TRIAD3A is reported to form K63 chains in response to phosphorylation^24^. K63 polyUb can activate TRIAD3A E3 ligase, but unlike other polyubiquitin chains, K63 chains inhibit TRIAD3A mediated LLPS (Figure 3c). This suggests that TRIAD3A autophagosome function may be regulated both by signaling/phosphorylation and by the local polyubiquitin environment. Interestingly, TRIAD3A is also enriched in the nucleus and our screen for TRIAD3A substrates identified multiple candidates that function in the nucleus including TDP43 and Fus. Like tau, several candidates form pathological aggregates in neurodegenerative diseases^3^. In addition to direct ubiquitination by TRIAD3A, it is possible that proteins are polyubiquitinated by other E3 ligases and that TRIAD3A, via its UBA domain, binds and redirects substrates to autophagosomes. This model can rationalize the observed pathological feature of ubiquitin accumulation and intranuclear inclusion in the nucleus of neurons in Gordon-Holms syndrome^20^. The distinct ubiquitin chain activating conditions for TRIAD3A versus SQSTM1/p62 and NBR1 anticipate dynamic and potentially reciprocal delivery of tau and other substrates to lysosome.

Tau aggregates accumulate and spread in the brain of individuals with neurodegenerative diseases, and a central unanswered question is how pathologic tau aggregation is initiated. Evidence here indicates that TRIAD3A induces tau fibrillization within the LLPS, and in conditions of normal lysosome function the TRIAD3A-autophagosome pathway effectively degrades tau and, for example, protects neurons from the pathological effects of Tau P301S. However, in the condition of reduced lysosomal function amyloid tau accumulates in association with TRIAD3A (Extended data Figure 6a and b). Several lines of evidence indicate that neurodegenerative diseases are closely linked to the disruption of lysosomal function ^6, 52,53^. Moreover, enlarged endosomes are evident in brains of individuals genetically predisposed to AD prior to cognitive decline suggesting that endolysosomal defects occur prior to clinically defined disease onset ^53^. Since TRIAD3A mediated tau homeostasis is dependent on lysosome function, the resulting disrupted function of TRIAD3A may increase fibrillar tau and thereby contribute to the initiation of pathological tau accumulation ^54-56^. In the AD brain, TRIAD3A’s contribution to tau homeostasis is further decreased by its down-regulated expression, and TRIAD3A-associated tau accumulates in various fibrillar forms consistent with a stall of the lysosome degradative process. The dependence of TRIAD3A selective autophagy on lysosome function and its capacity to induce tau fibrillation makes it a compelling target pathway in neurodegenerative pathophysiology.

## Acknowledgments

We thank Robert N. Cole at the Mass Spectrometry and Proteomics Facility in Johns Hopkins University and Ross Tomaino at Taplin Mass Spectrometry Facility in Harvard Medical School for mass spectrometry analysis. We thank the Janelia’s Virus Tool facility for helping prepare the AAV. Research support was provided by R01 MH053608 (P.F.W.). The Brain Resource Center is supported by the JHU Alzheimer’s Disease Research Center (NIH P30AG066507).

## Author contributions

W.Z. conceived and designed the project, performed *in vitro* experiments and interpreted results. Y.A generated the TRIAD3A-R694C mutation mice. A.P. made contribution to COIP experiment. J.R. A.B and J.T. provided the frontal cortex human tissues and help with the human brain slice staining and analysis. J.Z. performed all of other *in vivo* experiments and imaging analysis. All authors analyzed the data. W.Z., J.Z. and P.F.W wrote the manuscript with input from other authors. P.F.W and W.Z. supervised the work.

## Inclusion and diversity

We support inclusive, diverse, and equitable conduct of research.

## References

1. Spillantini, M.G. & Goedert, M. Tau protein pathology in neurodegenerative diseases. Trends in neurosciences 21, 428–433 (1998).

2. Ballatore, C., Lee, V.M. & Trojanowski, J.Q. Tau-mediated neurodegeneration in Alzheimer’s disease and related disorders. Nature reviews. Neuroscience 8, 663–672 (2007).

3. Higashi, S. et al. Concurrence of TDP-43, tau and alpha-synuclein pathology in brains of Alzheimer’s disease and dementia with Lewy bodies. Brain research 1184, 284–294 (2007).

4. Robinson, J.L. et al. The development and convergence of co-pathologies in Alzheimer’s disease. Brain 144, 953–962 (2021).

5. Schweighauser, M. et al. Age-dependent formation of TMEM106B amyloid filaments in human brains. Nature (2022).

6. Chang, A. et al. Homotypic fibrillization of TMEM106B across diverse neurodegenerative diseases. Cell 185, 1346–1355 e1315 (2022).

7. Jiang, Y.X. et al. Amyloid fibrils in disease FTLD-TDP are composed of TMEM106B not TDP-43. Nature (2022).

8. Xu, Y., Zhang, S. & Zheng, H. The cargo receptor SQSTM1 ameliorates neurofibrillary tangle pathology and spreading through selective targeting of pathological MAPT (microtubule associated protein tau). Autophagy 15, 583–598 (2019).

9. Ramesh Babu, J. et al. Genetic inactivation of p62 leads to accumulation of hyperphosphorylated tau and neurodegeneration. J Neurochem 106, 107–120 (2008).

10. Ma, X. et al. CCT2 is an aggrephagy receptor for clearance of solid protein aggregates. Cell 185, 1325–1345 e1322 (2022).

11. Darwich, N.F. et al. Autosomal dominant VCP hypomorph mutation impairs disaggregation of PHF-tau. Science 370 (2020).

12. Zhang, Z.Y. et al. TRIM11 protects against tauopathies and is down-regulated in Alzheimer’s disease. Science 381, eadd6696 (2023).

13. Lovestam, S. et al. Assembly of recombinant tau into filaments identical to those of Alzheimer’s disease and chronic traumatic encephalopathy. Elife 11 (2022).

14. Boyko, S., Surewicz, K. & Surewicz, W.K. Regulatory mechanisms of tau protein fibrillation under the conditions of liquid-liquid phase separation. Proc Natl Acad Sci U S A 117, 31882–31890 (2020).

15. Rai, S.K., Savastano, A., Singh, P., Mukhopadhyay, S. & Zweckstetter, M. Liquid-liquid phase separation of tau: From molecular biophysics to physiology and disease. Protein Sci 30, 1294–1314 (2021).

16. Babinchak, W.M. & Surewicz, W.K. Liquid-Liquid Phase Separation and Its Mechanistic Role in Pathological Protein Aggregation. J Mol Biol 432, 1910–1925 (2020).

17. Wegmann, S. et al. Tau protein liquid-liquid phase separation can initiate tau aggregation. EMBO J 37 (2018).

18. Spratt, D.E., Walden, H. & Shaw, G.S. RBR E3 ubiquitin ligases: new structures, new insights, new questions. Biochem J 458, 421–437 (2014).

19. Santens, P. et al. RNF216 mutations as a novel cause of autosomal recessive Huntington-like disorder. Neurology 84, 1760–1766 (2015).

20. Margolin, D.H. et al. Ataxia, dementia, and hypogonadotropism caused by disordered ubiquitination. N Engl J Med 368, 1992–2003 (2013).

21. Alqwaifly, M. & Bohlega, S. Ataxia and Hypogonadotropic Hypogonadism with Intrafamilial Variability Caused by RNF216 Mutation. Neurol Int 8, 6444 (2016).

22. Calandra, C.R. et al. Gordon Holmes Syndrome Caused by RNF216 Novel Mutation in 2 Argentinean Siblings. Mov Disord Clin Pract 6, 259–262 (2019).

23. Chen, K.L. et al. Whole-Exome Sequencing Identified a Novel Mutation in RNF216 in a Family with Gordon Holmes Syndrome. J Mol Neurosci 72, 691–694 (2022).

24. Cotton, T.R. et al. Structural basis of K63-ubiquitin chain formation by the Gordon-Holmes syndrome RBR E3 ubiquitin ligase RNF216. Molecular cell 82, 598–615 e598 (2022).

25. Mabb, A.M. et al. Triad3A regulates synaptic strength by ubiquitination of Arc. Neuron 82, 1299–1316 (2014).

26. Schwintzer, L., Aguado Roca, E. & Broemer, M. TRIAD3/RNF216 E3 ligase specifically synthesises K63-linked ubiquitin chains and is inactivated by mutations associated with Gordon Holmes syndrome. Cell Death Discov 5, 75 (2019).

27. Seenivasan, R. et al. Mechanism and chain specificity of RNF216/TRIAD3, the ubiquitin ligase mutated in Gordon Holmes syndrome. Human molecular genetics 28, 2862–2873 (2019).

28. Marin, I. RBR ubiquitin ligases: Diversification and streamlining in animal lineages. J Mol Evol 69, 54–64 (2009).

29. van Wijk, S.J. & Timmers, H.T. The family of ubiquitin-conjugating enzymes (E2s): deciding between life and death of proteins. FASEB journal : official publication of the Federation of American Societies for Experimental Biology 24, 981–993 (2010).

30. Swatek, K.N. et al. Insights into ubiquitin chain architecture using Ub-clipping. Nature 572, 533–537 (2019).

31. Scott, D., Oldham, N.J., Strachan, J., Searle, M.S. & Layfield, R. Ubiquitin-binding domains: mechanisms of ubiquitin recognition and use as tools to investigate ubiquitin-modified proteomes. Proteomics 15, 844–861 (2015).

32. Hicke, L., Schubert, H.L. & Hill, C.P. Ubiquitin-binding domains. Nat Rev Mol Cell Biol 6, 610–621 (2005).

33. Shin, Y. & Brangwynne, C.P. Liquid phase condensation in cell physiology and disease. Science 357 (2017).

34. Hahn, S. Phase Separation, Protein Disorder, and Enhancer Function. Cell 175, 1723–1725 (2018).

35. Wang, Z. & Zhang, H. Phase Separation, Transition, and Autophagic Degradation of Proteins in Development and Pathogenesis. Trends Cell Biol 29, 417–427 (2019).

36. Noda, N.N., Wang, Z. & Zhang, H. Liquid-liquid phase separation in autophagy. The Journal of cell biology 219 (2020).

37. Gatica, D., Lahiri, V. & Klionsky, D.J. Cargo recognition and degradation by selective autophagy. Nature cell biology 20, 233–242 (2018).

38. Kirkin, V. & Rogov, V.V. A Diversity of Selective Autophagy Receptors Determines the Specificity of the Autophagy Pathway. Molecular cell 76, 268–285 (2019).

39. Jumper, J. et al. Highly accurate protein structure prediction with AlphaFold. Nature 596, 583–589 (2021).

40. Jatana, N., Ascher, D.B., Pires, D.E.V., Gokhale, R.S. & Thukral, L. Human LC3 and GABARAP subfamily members achieve functional specificity via specific structural modulations. Autophagy 16, 239–255 (2020).

41. Bhutia, S.K. et al. Monitoring and Measuring Mammalian Autophagy. Methods in molecular biology (Clifton, N.J.) 1854, 209–222 (2019).

42. Dooley, H.C. et al. WIPI2 links LC3 conjugation with PI3P, autophagosome formation, and pathogen clearance by recruiting Atg12-5-16L1. Molecular cell 55, 238–252 (2014).

43. Branon, T.C. et al. Efficient proximity labeling in living cells and organisms with TurboID. Nat Biotechnol 36, 880–887 (2018).

44. Guo, Q. et al. Targeted Quantification of Detergent-Insoluble RNA-Binding Proteins in Human Brain Reveals Stage and Disease Specific Co-aggregation in Alzheimer’s Disease. Frontiers in molecular neuroscience 14, 623659 (2021).

45. Schmued, L. et al. Introducing Amylo-Glo, a novel fluorescent amyloid specific histochemical tracer especially suited for multiple labeling and large scale quantification studies. J Neurosci Methods 209, 120–126 (2012).

46. Kaminskyy, V., Abdi, A. & Zhivotovsky, B. A quantitative assay for the monitoring of autophagosome accumulation in different phases of the cell cycle. Autophagy 7, 83–90 (2011).

47. Vinod, V., Padmakrishnan, C.J., Vijayan, B. & Gopala, S. ‘How can I halt thee?’ The puzzles involved in autophagic inhibition. Pharmacol Res 82, 1–8 (2014).

48. Wegmann, S. et al. Experimental evidence for the age dependence of tau protein spread in the brain. Science Advances 5, eaaw6404 (2019).

49. Grumati, P. & Dikic, I. Ubiquitin signaling and autophagy. The Journal of biological chemistry 293, 5404–5413 (2018).

50. Agudo-Canalejo, J. et al. Wetting regulates autophagy of phase-separated compartments and the cytosol. Nature 591, 142–146 (2021).

51. Mathieu, C., Pappu, R.V. & Taylor, J.P. Beyond aggregation: Pathological phase transitions in neurodegenerative disease. Science 370, 56–60 (2020).

52. Van Acker, Z.P., Bretou, M. & Annaert, W. Endo-lysosomal dysregulations and late-onset Alzheimer’s disease: impact of genetic risk factors. Molecular neurodegeneration 14, 20 (2019).

53. Lee, J.H. et al. Faulty autolysosome acidification in Alzheimer’s disease mouse models induces autophagic build-up of Abeta in neurons, yielding senile plaques. Nat Neurosci 25, 688–701 (2022).

54. Arakhamia, T. et al. Posttranslational Modifications Mediate the Structural Diversity of Tauopathy Strains. Cell 180, 633–644 e612 (2020).

55. Kametani, F. et al. Comparison of Common and Disease-Specific Post-translational Modifications of Pathological Tau Associated With a Wide Range of Tauopathies. Front Neurosci 14, 581936 (2020).

56. Wesseling, H. et al. Tau PTM Profiles Identify Patient Heterogeneity and Stages of Alzheimer’s Disease. Cell 183, 1699–1713 e1613 (2020).

